# Tissue resident cells differentiate *S. aureus* from *S. epidermidis* via IL-1ß following barrier disruption in healthy human skin

**DOI:** 10.1101/2024.02.19.580932

**Authors:** Julia C. Lang, Andreas Brutscher, Marcus Ehrström, Keira Melican

## Abstract

The *Staphylococcus* species are a dominant part of the human skin microbiome and present across all body sites. *Staphylococcus epidermidis* is a ubiquitous skin commensal, while *S. aureus* is thought to colonize at least 30% of the population. *S. aureus* are not only colonizers but a leading cause of skin and soft tissue infections and a critical healthcare concern. To understand how healthy human skin may differentiate commensal bacteria, such as *S. epidermidis*, from the potential pathogen methicillin-resistant *S. aureus* (MRSA),, we use *ex vivo* human skin models that allows us to study this host-bacterial interaction in the most clinically relevant environment. Our work highlights the role of the outer stratum corneum as a protective physical barrier against invasion by colonizing Staphylococci. We show how the structural cells of the skin can internalize and respond to different Staphylococci with increasing sensitivity. Disruption of the protective stratum corneum triggered an increased and discriminative immune response led by strong IL-1ß expression. We identified and localized tissue resident Langerhans cells (LCs) as a potential source of this IL-1ß and go on to show a dose dependent response of MUTZ-LCs to *S. aureus* but not *S. epidermidis*. This suggests an important role of LCs in sensing and discriminating between bacteria in healthy human skin. This work provides a detailed snapshot of how human skin differentiates between friend and potential foe. With the rise in antibiotic resistance, understanding the innate immune response of healthy skin may help us find ways to enhance or manipulate these natural defenses to help prevent invasive infection.

**Author Summary:** How healthy human skin can differentiate between commensal bacteria which live harmoniously on our skin and those bacteria which have the potential to cause invasive infection is of great interest. Here we use *ex vivo* human skin models to show how the outer layers of human skin resists bacterial invasion and that disruption of this layer leads to increased infection and inflammation. We show how colonization of disrupted skin by the potentially pathogenic *Staphyloccus aureus* causes an increased IL-1ß signaling compared to the commensal *S. epidermidis*. We show that tissue resident Langerhans cells are well positioned throughout the skin epidermis and can differentiate commensal from pathogen via IL-1ß. This work identifies one of the subtle, yet effective pathways by which human skin differentiates commensals and pathogens. Understanding these mechanisms could lead to potential new treatment and prevention targets, mechanisms by which we can enhance innate immunity.

## Introduction

Human skin, our largest organ, forms the first physical and immunological defense against bacteria. Paradoxically, our skin is also covered by a highly diverse and dynamic microbiome which is believed to play a direct role in protecting us from invasive bacterial pathogens(1). The *Staphylococcus* species are a prominent feature of the human skin microbiome, with *S. epidermidis*, the most common coagulase negative Staphylococci (CoNS), comprising over 90% of the skin resident aerobic flora(2). Around 20% of the human population are also colonized with the potentially pathogenic *S. aureus*, with another 30% being transient carriers(3). *S. aureus* is responsible for 76% of all skin and soft tissue infections and numerous invasive infections(4,5). Methicillin resistant *S. aureus* (MRSA) causes the highest proportion of deaths related to antibiotic resistant bacteria in the developed world(6,7). An ongoing challenge in research is to understand how these bacteria can be both a harmless commensal and a potentially deadly pathogen. In this work we sought to understand how healthy human skin recognizes and responds to colonizing *Staphylococcus*, and how skin may differentiate between *S. epidermidis* and *S. aureus*. Understanding how our skin protects us in the vast majority of cases we come into contact with *S. aureus* will give greater clarity to our first line of defense and open the possibilities for enhancing or manipulating our innate protection when needed.

For many years, studying host-pathogen interaction between *Staphylococcus sp.* and skin has been performed using mouse models(8–10). While adding great knowledge to the field, it is becoming increasingly evident that the human skin response to *Staphylococcus* differs significantly from mouse. Mice are not natural hosts for human *Staphylococcus* isolates(11) and the skin of rodents and humans have dramatically different structure and immunological function(12). Additionally, elements of *Staphylococcus* pathogenesis have been shown to be human specific, including the interaction of Staphylococcal toxins with human neutrophils, and between *Staphylococcus* and human langerin, the receptor of skin resident Langerhans cells(13,14). These factors highlight the importance of studying host-pathogen interactions in the most clinically relevant microenvironment, human skin(15,16). In our group, we have developed several human skin models to address host-pathogen interaction. We previously published work demonstrating that in a xenografted model, where healthy human skin is grafted onto immunocompromised mice, the bacterial colonization patterns and inflammatory response of human skin to MRSA colonization differed significantly from previous mouse models(17). In this current work, we have optimized explant human skin models to differentiate the response of healthy human skin to *S. epidermidis* vs MRSA. We combine this with cell culture to understand the role of each cell type in this early response. An advantage of our adult human skin models is that they contain the full complement of tissue resident immune cells and here we focus on Langerhans cells (LCs) and their ability to differentiate *S. epidermidis* vs MRSA. These skin resident immune cells are often described as a bridging element between innate and adaptive immunity and play an important role in activating specific T-cell responses(18).

Our results reveal the critical role of the human skin outer layer, the stratum corneum, in providing a physical barrier to bacterial invasion. We also demonstrate how the skin structural cells can internalize and respond to bacteria, with an increasing responsiveness as we move into deeper layers of the skin. We then delve further into the different inflammatory response to MRSA and *S. epidermidis* and identify Langerhans cells as a potential driver of this differentiation. This work offers a high resolution understanding of the early phases of human skin colonization and how our skin differentiates friend from foe.

## Results

### An intact stratum corneum forms the first line of defense against *Staphylococcus sp*

We previously showed in an *in vivo* xenograft model that the human stratum corneum is an important barrier to MRSA invasion(17). In this work, we studied the importance of an intact stratum corneum during colonization by commensal vs pathogenic bacteria. To do this we optimized an *ex vivo* tissue explant model using biopsies of fresh, healthy, human skin from multiple donors. The integrity and morphology of the skin model was validated after 72 h of *ex vivo* culture (Fig S1). These full thickness biopsies contain an intact stratum corneum atop the epidermal and dermal layers. The surface of the skin was topically inoculated with ∼1.6×10^6^ CFU/cm^2^ *S. aureus* USA300 (MRSA)(19), or *S. epidermidis* 1457(20). This inoculation dose was correlated with the reported density of the human skin microbiota(21). Following 48 h of colonization, there was no significant differences in the ability of *S. epidermidis* or MRSA to colonize the skin, as measured by CFU, though there was a trend towards higher MRSA CFUs (Fig 1A). Individual variability between donors was seen. In our xenograft model, we had shown an IL-8 response to skin colonization with MRSA(17) and we therefore tested for potential inflammatory differences between colonization by MRSA or *S. epidermidis*. No significant difference in either IL-8 (Fig 1B) or CXCL12 (Fig 1C) concentrations were seen between colonization with MRSA or *S. epidermidis*. The time frames of this *ex vivo* model were significantly shorter than those reported for the xenograft and we therefore evaluated the localization of bacteria at this 48 h timepoint. Confocal microscopy revealed that the majority of *S. aureus* USA300 expressing GFP were found adhered to the intact stratum corneum (Fig 1D). This is of importance as this outer layer is lacking from mouse models and often tape stripped away in other human models(22–24). No significant bacterial penetration into the tissue was seen, correlating to our previous work with the xenograft(17). We did however locate occasional intracellular bacteria in nucleated cells of the upper epidermal layers (Fig 1E). This data reinforced the important role of the stratum corneum as the key initial binding site for *Staphylococcus sp*. on the skin, and the integrity of the skin barrier to protect against tissue invasion but also suggested a role for intracellular bacteria during skin colonization.

**Fig 1.**
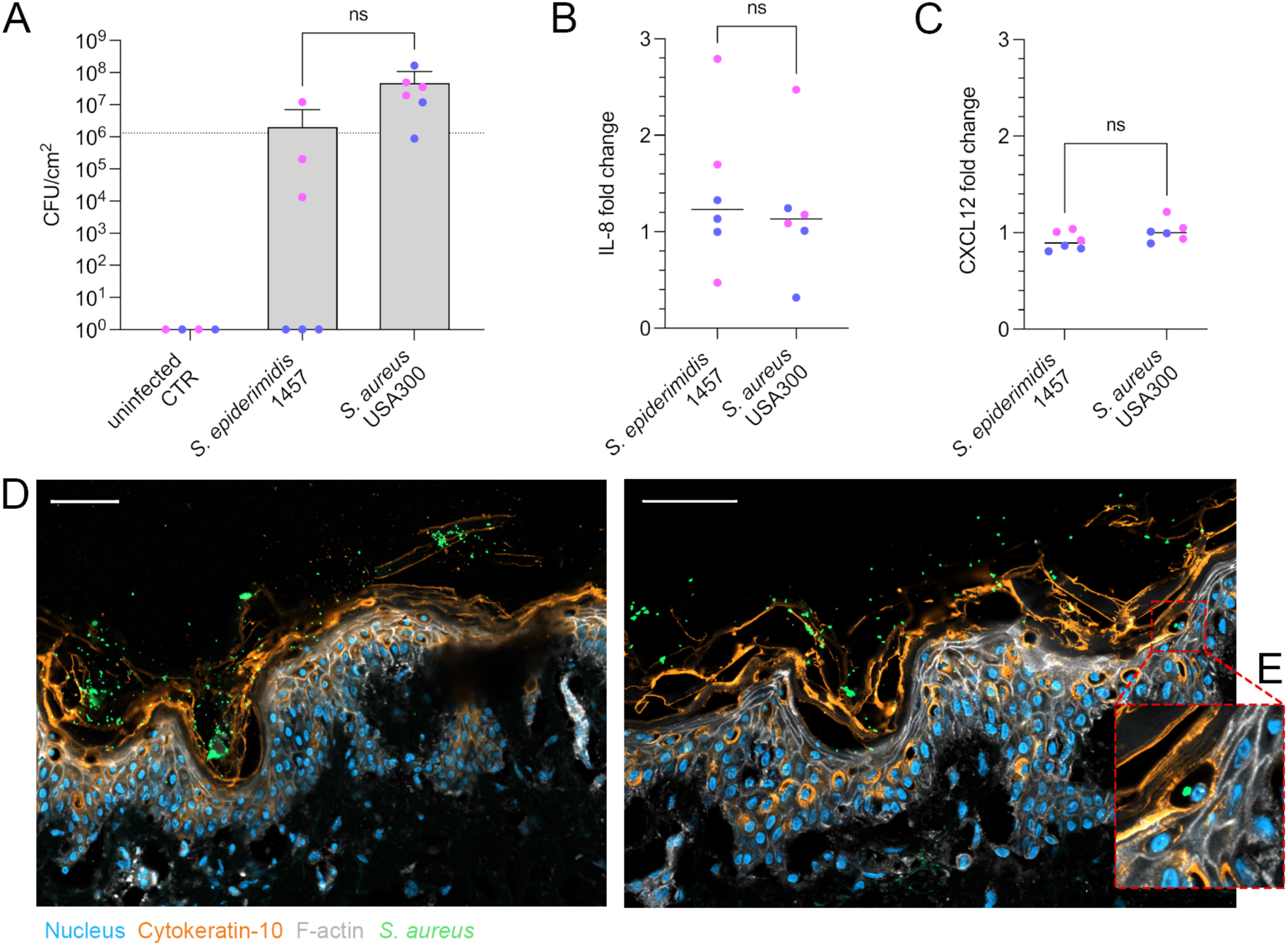
Inflammatory response to colonizing *Staphylococcus sp.* in human skin. (**A**) CFU counts of *S. epidermidis* 1457 and *S. aureus* USA300 48 h after epicutaneous colonization of *ex vivo* human skin. Data show means ± SD. Dots represent individual data points, colours differentiate skin donors (n=6). Dotted line depicts inoculum. Significance determined by one-way ANOVA with Tukey’s correction for multiple comparisons. (**B-C**) ELISA IL-8 (**B**) and CXCL12 (**C**), as x-fold change over uninfected controls. Dots represent individual data points, colours differentiate skin donors (n=6). Significance determined by unpaired t-test with Welch’s correction. (**D**) Representative immunofluorescence staining of cytokeratin-10 (orange), f-actin (white), nuclei (blue) in skin model topically colonized with *S. aureus* USA300 GFP (green) for 48 h. Scale bars = 50µm. N=3. (**E**) Insert of area in (**D**) showing intracellular *S. aureus* USA300 GFP.

### Staphylococci can be internalized by skin structural cells with differential inflammatory responses

The intracellular localization of *Staphylococcus* may be a means to penetrate into deeper tissue, but it has also been suggested that intracellular localization plays a role in chronic wounds and recurrent infection(25–27). To investigate the role of intracellular bacteria in the human skin response to colonization we studied the ability of *S. epidermidis* and MRSA to internalize in the key structural cells of the human skin. We included primary isolated human corneocytes, human keratinocytes (HEKa) and dermal fibroblasts (HDFn). We challenged the cells with an MOI of 1:100 of both *S. epidermidis* and MRSA for 3 h before gentamicin-lysostaphin treatment. The cells were further incubated until 24 h p.i. Intracellular CFU was determined after 3 h (Fig 2A) and IL-8 expression after 24 h (Fig 2B). Corneocytes demonstrated no intracellular uptake or invasion of either *S. epidermidis* or MRSA and no expression of IL-8 (Fig 2A, B). In line with their abilities and functions as non-professional phagocytes, keratinocytes and fibroblasts harbored intracellular bacteria, both *S. epidermidis* and MRSA (Fig 2A). Keratinocytes contained similar numbers of *S. epidermidis* and MRSA while fibroblasts contained significantly more MRSA than *S. epidermidis*. The MRSA numbers in the fibroblasts were significantly higher than those found in keratinocytes (Fig 2A). Keratinocytes displayed a low IL-8 response to both *S. epidermidis* and MRSA while fibroblasts displayed higher IL-8 responses and a significantly higher response to MRSA as compared to *S. epidermidis* (Fig 2B). Microscopy analysis of cells infected with *S. aureus* GFP for 3 h validated these results indicating no intracellular uptake of MRSA into corneocytes, while keratinocytes and fibroblasts showed distinct patterns for intracellular MRSA (Fig 2C). The keratinocytes infection was heterogenous with individual cells harboring numerous clustered MRSA. Fibroblasts were more homogenously infected with intracellular bacteria spreading throughout the cell body (Fig 2C).

**Fig 2.**
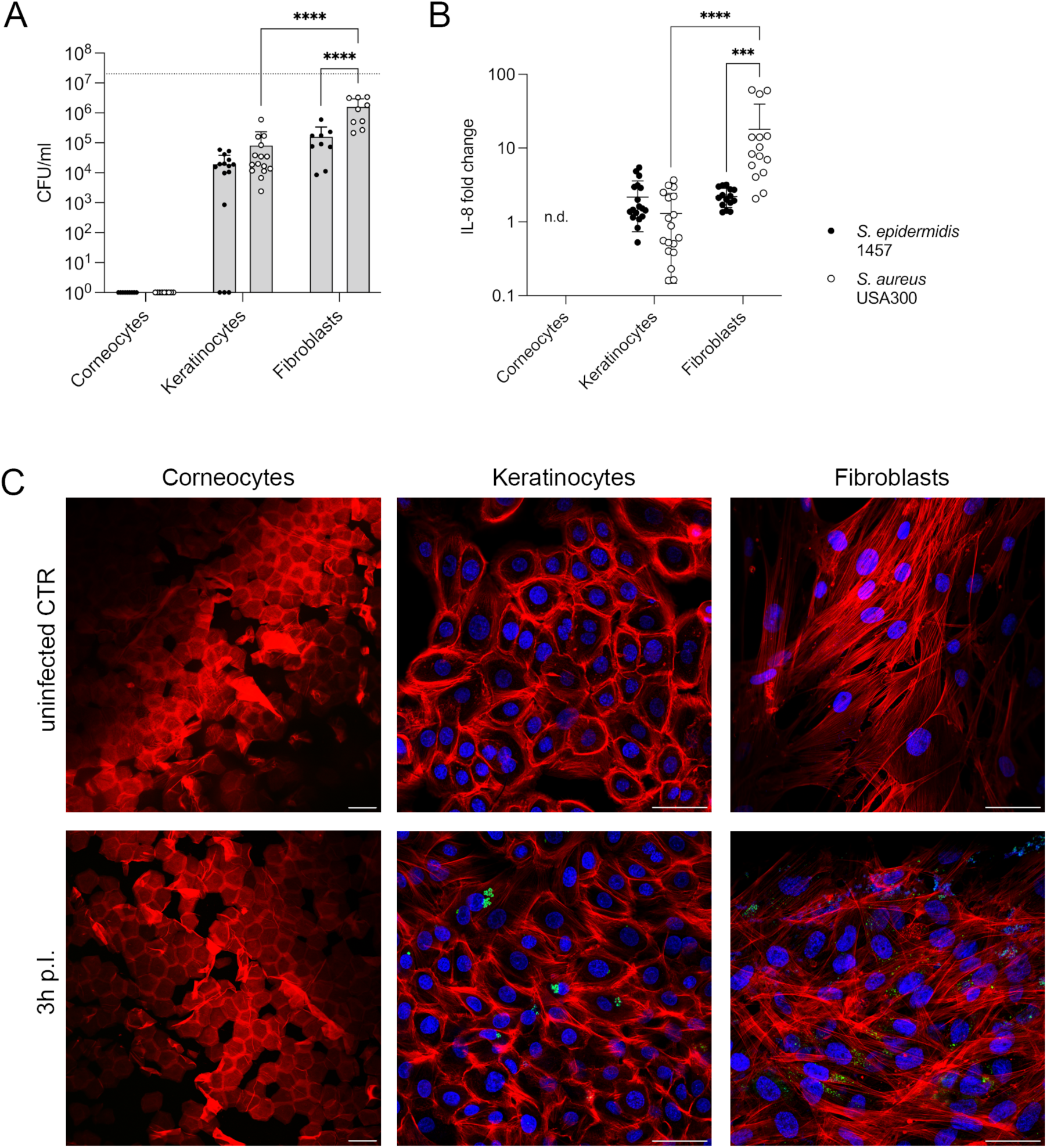
Intracellular Staphylococcus sp. induce an IL-8 response in keratinocytes and fibroblasts, but not corneocytes. (**A**) Intracellular CFU counts of *S. epidermidis* 1457 (filled circles) and *S. aureus* USA300 (empty circles) 3 h p.i. of tape-stripped corneocytes, human keratinocytes (HEKa) and fibroblasts (HDFn). Bars represent means ± SD, dots show individual data points (corneocytes and fibroblasts n=3; keratinocytes n=5; 3 technical replicates). Dotted line = initial inoculum. Significance was determined by a two-way ANOVA with Tukey’s correction for multiple comparisons. Significance is denoted by **** p ≤ 0.0001. (**B**) IL-8 response displayed as x-fold change over uninfected controls measured by ELISA in cell culture supernatants of *S. epidermidis* 1457 or *S. aureus* USA300 infected corneocytes, keratinocytes or fibroblasts. Dot plot shows individual data points (corneocytes and fibroblasts n = 3; keratinocytes n = 5; 3 technical replicates each). Significance was determined by a two-way ANOVA with Tukey’s correction for multiple comparisons. Significance is denoted by *** p ≤ 0.001 or **** p ≤ 0.0001. (**C**) Microscopic detection of intracellular *S. aureus* GFP in cell monolayers of tape-stripped corneocytes, human keratinocytes (HEKa) or fibroblasts (HDFn); uninfected CTR vs 3h infection. Immunostaining of f-actin (red) and nuclei (blue) in HEKa and HDFn, as well as autofluorescence staining of corneocytes (red). Scale bars = 50µm.

This data demonstrates that while corneocytes are the key binding target in early skin colonization, these cells do not internalize *Staphylococcus* bacteria or initiate an apparent inflammatory response. Both keratinocytes and fibroblasts could internalize *Staphylococcus* and generate an IL-8 inflammatory signal, with fibroblasts demonstrating greater inflammatory and particularly discriminatory potential between *S. epidermidis* and MRSA. This suggests a striated tissue response from the structural cells graduating from non-responsive corneocytes on the outer skin surface through the keratinocyte rich epidermis to the most responsive fibroblasts located in the skin dermis.

### Disruption of the stratum corneum enhances *S. aureus* colonization and tissue inflammatory response

This data highlighted differences in the skin layers, with the outer most layers being non-responsive to *Staphylococcus*, but the structural cells located in the deeper tissue more capable of discriminating MRSA from *S. epidermidis*. Recent, ground breaking, work has shown that MRSA, via V8 protease, leads to an urge to itch skin(28). Scratching leads to minor disruptions in the outer layers of skin and we wanted to study if these minor disruptions would trigger a more substantial inflammatory skin response. We altered our tissue model by superficially scratching the surface of the human skin before bacterial colonization for 48 h (Fig S2). This barrier disruption led to significantly higher loads of MRSA compared to *S. epidermidis*, with noted variation between individual skin donors (Fig 3A). Inoculation of scratched skin with MRSA led to significantly higher expression of IL-8 than inoculation with *S. epidermidis* (Fig 3B). Scratched skin inoculated with either MRSA or *S. epidermidis* had increased IL-8 compared to their donor matched non-infected controls and for MRSA also a significant increase compared to unscratched skin (Fig 3B, 1B). Expression of CXCL12 was however not significantly altered, indicating a specificity in this response (Fig 3C). This data further supports the role of the stratum corneum in protecting the tissue, showing how minor disruptions in this layer leads to increasing bacterial loads as well as inflammatory signaling, particularly for MRSA.

**Fig 3.**
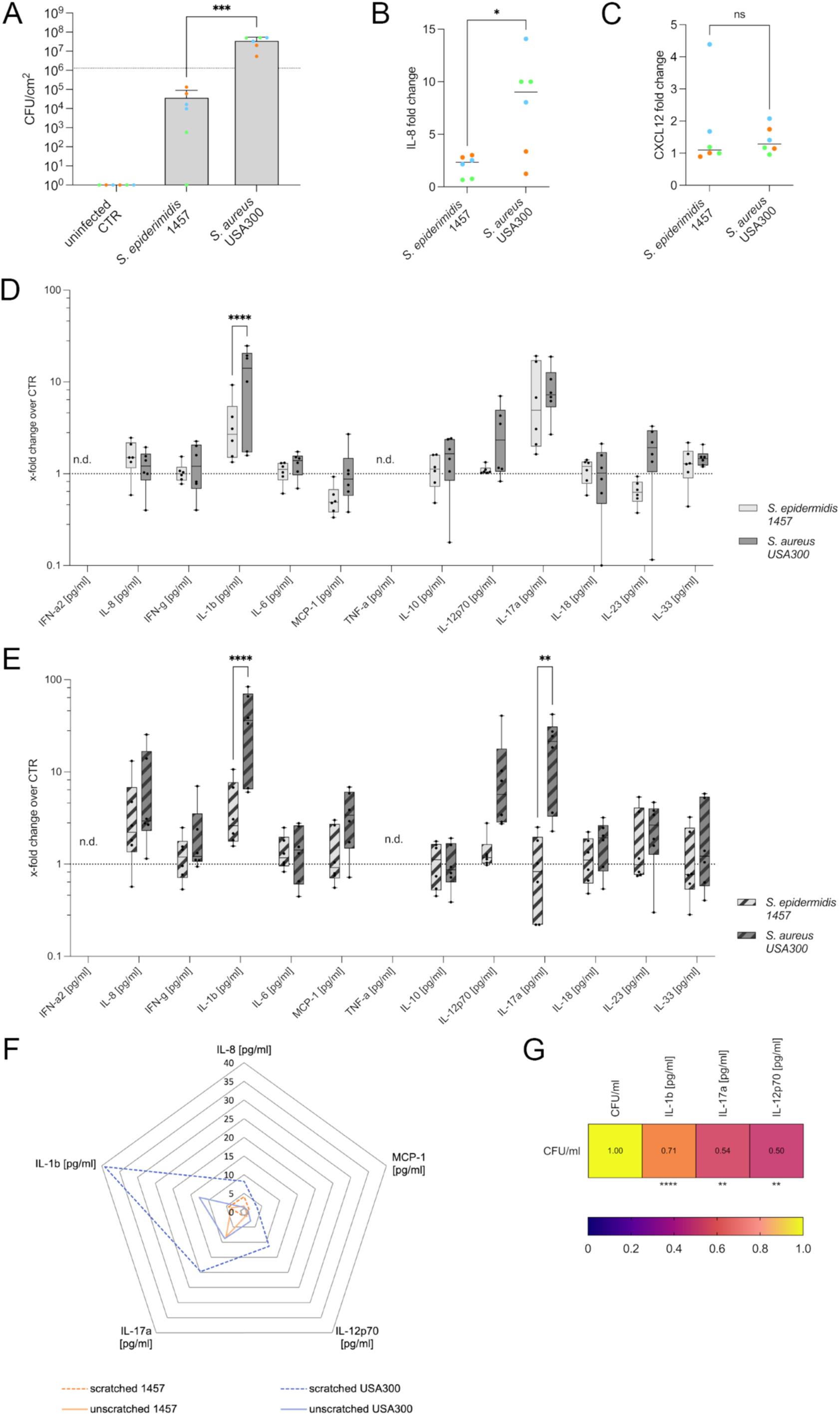
Stratum corneum disruption results in increased inflammatory response to *S. aureus* colonization. (**A**) CFU counts of *S. epidermidis* 1457 and *S. aureus* USA300 48 h after epicutaneous colonization of biopsies with disrupted stratum corneum (s.c.). Bars = means ± SD, dots show individual data points (n=6, skin donors indicated by colors). Dotted line depicts inoculum. Significance determined by one-way ANOVA with Tukey’s correction for multiple comparisons. Significance denoted by *** p ≤ 0.001. (**B**) IL-8 and (**C**) CXCL12 response displayed as x-fold change over uninfected controls measured by ELISA in homogenates of skin biopsies with disrupted s.c. inoculated with *S. epidermidis* 1457 or *S. aureus* USA300. Dot plot shows individual data points (n=6, skin donors indicated by colors). Significance determined by unpaired t test with Welch’s correction. Significance denoted by * p ≤ 0.05. (**D-E**) Cytokine expression displayed as x-fold change from uninfected controls in colonized skin explants with an intact s.c. (**D**) or disrupted s.c. (**E**) measured by LEGENDplex™ bead-based immunoassay. The box plots represent interquartile range (IQR) with the median shown as black line. Dots show individual data points (n=6, 2-3 individual skin donors). Statistical significance determined by two-way ANOVA with Šídák’s correction for multiple comparisons and denoted by ** p ≤ 0. 01 and **** p ≤ 0. 0001. (**F**) Selected cytokines on a radar plot shown as mean x-fold changes over uninfected controls. (**G**) Spearman correlation between CFUs and IL-1β, IL-17A and IL-12p70 measured in human skin biopsies 48 h p.i. Significance for each cytokine compared to the CFU/ml denoted by ** p ≤ 0. 01 and **** p ≤ 0. 0001.

To further evaluate the inflammatory differences in scratched and unscratched skin we expanded our analysis to a panel of inflammatory cytokines including IL-1β, IFN-α2, IFN-γ, TNF-α, MCP-1, IL-6, CXCL8 (IL-8), IL-10, IL-12p70, IL-17A, IL-18, IL-23 and IL-33. In intact, unscratched skin, only IL-17A and IL-1β were significantly increased above the non-infected controls (Fig 3D). Interestingly IL-1β expression was significantly higher in MRSA inoculated tissue compared to *S. epidermidis* (Fig 3D). In tissues with a disrupted, scratched stratum corneum, the cytokine responses were increased for most factors (Fig 3E). Both IL-1β and IL-17A showed significantly different expression when colonized by MRSA compared to *S. epidermidis* (Fig 3E). For better visualization, we created a radar plot of the mean x-fold changes for the most significantly upregulated factors, IL-1β, MCP-1, CXCL8 (IL-8), IL-12p70 and IL-17A (Fig 3F). This showed the IL-1β, IL-17A and IL-12p70 dominance of the skin response to MRSA which is enhanced 30x in the case of IL-1β following barrier disruption. A significant positive correlation between bacterial CFUs and IL-1β, IL-17A and IL-12p70 concentrations in individual biopsy was found (Fig 3G). IL-17A has long been reported to have an important role in cutaneous and mucosal host defense to control the microbiota and was found to be involved in defense against MRSA skin infection in mice. IL-12p70 is crucial to polarize naïve T cells towards a Th1 phenotype, but also has been described to induce Tregs and other immunosuppressive cells(33–36). IL-1β, the most strongly upregulated cytokine, has been described to promote neutrophil recruitment(10,37) and to modulate the expression of tight junction proteins(38).

This data demonstrates immunological differences in how healthy human skin responds to MRSA and *S. epidermidis* and the significant effect of barrier dysfunction on the amplitude of these responses. A key responsive factor identified in the human skin was IL-1β.

### Skin resident Langerhans cells do not internalize *Staphylococcus* but differentiate *S. aureus* from *S. epidermidis* via IL-1β

We then sought to investigate which cell types may be involved in the IL-1β response in both intact and scratched skin. Many cell types have been implicated in IL-1β signaling including keratinocytes, mast cell and Langerhans cells (LCs)(39–41). Our human skin model contains the full repertoire of tissue resident immune cells including LCs. To localize LCs in the tissue we performed confocal imaging of separated epidermal sheets. We identified the impressive phenotype of the LCs in the epidermis labelled with langerin (CD207) (Fig 4A). The cell bodies were located close to the epidermal, dermal basal membrane (Fig 4A, C), while the cell projections reached the outer surface of the skin (Fig 4B). This localization suggested a possible role for these cells in early identification of bacteria in both intact and disrupted skin. To study the individual role of the LCs in our tissue response to *S. epidermidis* and MRSA we differentiated a MUTZ-3 cell line into a LC phenotype. Successful differentiation of the cells was confirmed by analysis of langerin, CD1a and HLA-DR expression (Fig S3). We visually confirmed langerin expression and dendritic like morphology of the differentiated cells as previously described(42,43) (Fig 4D). To study the ability of the MUTZ-LCs to internalize and respond to MRSA or *S. epidermidis* we infected these cells with both bacteria at MOIs of 1,2,10 or 50. After 3 h of exposure we treated with gentamicin-lysostaphin. The cells were then incubated further to 24 h. No intracellular CFUs were found after 3 h, demonstrating a limited capacity for MUTZ-LCs to internalize Staphylococci. Despite the lack of bacterial uptake, we saw a significant, dose dependent expression of IL-1β in the cell culture media 24 h after MRSA exposure (Fig 4E). Exposure to *S. epidermidis* resulted in no IL-1β expression (Fig 4E). This data indicated than the IL-1β expression patterns we saw in the human skin correlates to MUTZ-LC exposure. IL-1β was the only cytokine differentially upregulated in intact skin exposed to MRSA vs *S. epidermidis,* which could be explained by the Langerhans cells projections at the skin surface. Scratched skin, in which there would be increased exposure, correlated to the higher IL-1β and significant differentiation between MRSA and *S. epidermidis*. This data suggests that LCs can differentiate MRSA from *S. epidermidis* and that these cells could play a key role in the IL-1β response we identified *in situ* in our human skin model.

**Fig 4.**
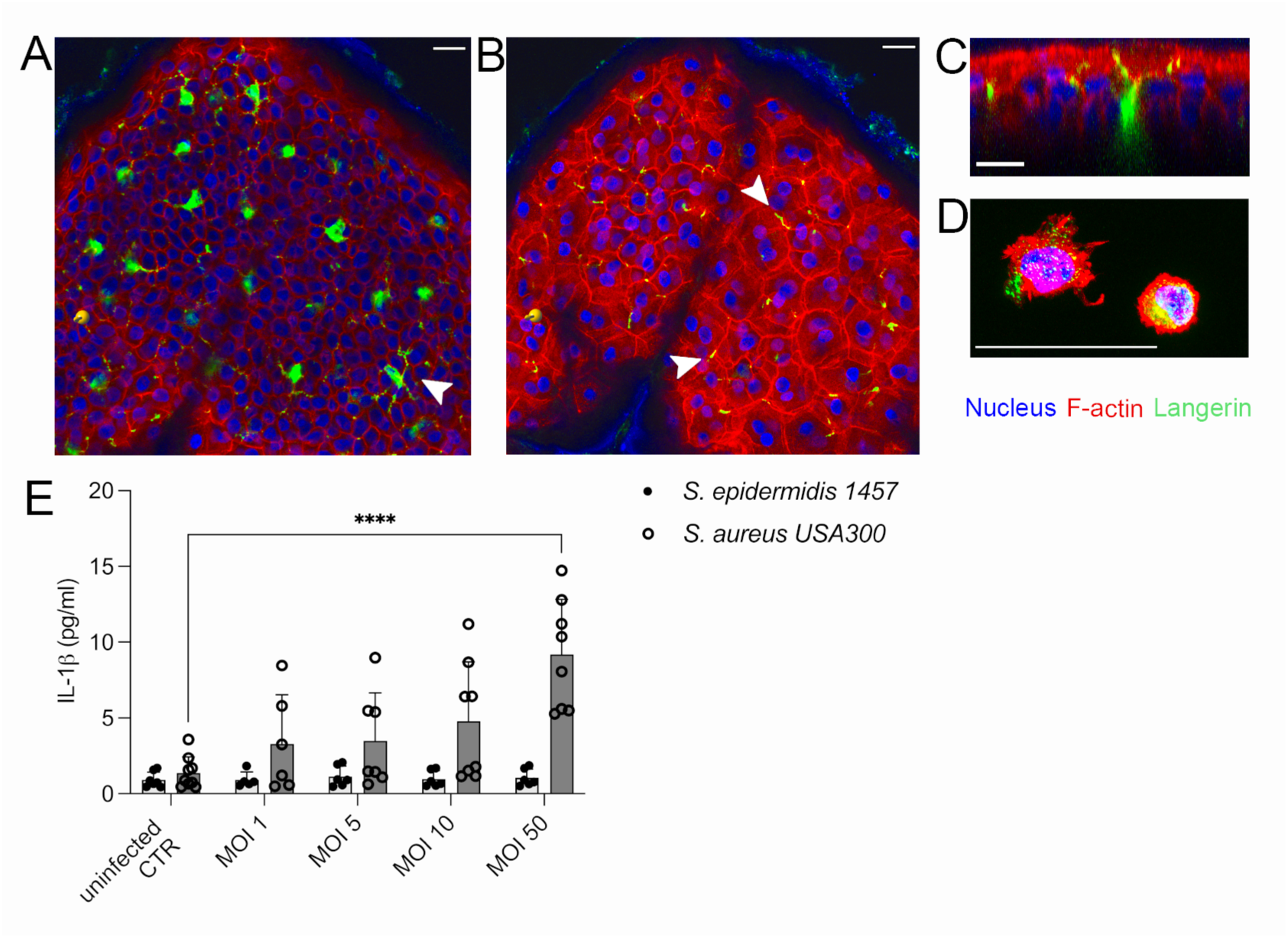
Skin resident Langerhans cells are located throughout the epidermis and can differentiate *S. aureus* from *S. epidermidis.* (**A**) Orthogonal projection of an epidermal sheet close to the basal membrane. Immunofluorescent staining of f-actin (red), nuclei (blue) and langerin (green). Scale bar = 20µm. Arrow indicates Langerhans cell (green) (**B**) Orthogonal projection of an epidermal sheet close to the stratum corneum. Immunofluorescent staining of f-actin (red), nuclei (blue) and langerin (green). Scale bar = 20µm. Arrows indicate overlap of Langerhans cells projections with keratinocytes cell walls. Scale bar = 20µm. (**C**) XZ projection of an epidermal sheet showing a Langerhans cell body at the base of the epidermis with projections extending to the skin surface. F-actin (red), nuclei (blue) and langerin (green). Scale bar = 20µm. (**D**) Maximum projection of MUTZ-LCs. F-actin (red), nuclei (blue) and langerin (green). Scale bar = 25µm. (**E**) IL-1β measured by ELISA in MUTZ-LCs supernatants at 24h p.i. after 3 h infection with *S. aureus* or *S. epidermidis*. Bars represent means ± SD, dots show individual data points (n=3-4 experimental repeats, with 2 technical replicates). Statistical significance determined by two-way ANOVA with Tukey’s correction for multiple comparisons. Statistical significance denoted by **** p ≤ 0.0001.

## Discussion

Our work in healthy human skin models advances our understanding of the first line of the human skin immune response to colonizing Staphylococci. We highlight the critical role of the stratum corneum in the initial protection to bacteria and how even minor disruption of this layer increase the bacterial ability to colonize and induce an inflammatory response. We show a potential role for intracellular bacteria during skin colonization and how the skin structural cells have differing sensitivities to staphylococci. We identify IL-1β in both scratched and unscratched skin and show how it is a key differentiator in the human skin response to the commensal *S. epidermidis* compared to the potentially pathogenic MRSA. Our data suggests tissue resident Langerhans cells are a major source of the IL-1β and demonstrates their ability to differentiate MRSA from *S. epidermidis*.

The new connections between epicutaneous *S. aureus* and itch shown by the Chiu lab(28) further highlights the relevance of our models as we seek to understand the early phases of host-bacterial interaction. While colonization of an intact stratum corneum showed very little immune response in the skin, transient surface colonization by MRSA may induce itch and the following skin damage in turn opens a window for the bacteria to infect deeper layers of the skin but also trigger a more pronounced immune response.

We demonstrate that the corneocytes in the stratum corneum do not internalize Staphylococci, but the underlying non-professional phagocytic keratinocytes and fibroblast cells can become intracellularly infected and have an increasing, graduated responsiveness. This finding is in line with others, showing that keratinocytes in stratum granulosum and stratum spinosum but also fibroblasts can take up bacteria or get invaded by them(1,29). This intracellular niche provides a hiding place from the host immune defense as well as many antibiotics(44) but both cell types have intracellular defense mechanisms to eliminate *S. aureus*(45–47). Skin diseases such as atopic dermatitis, commonly associated with compromised skin barrier, diabetic foot ulcers (DFUs), chronic non-healing wounds and recurrent infections, are all associated with intracellular *S. aureus*(48–52). Our work shows that intracellular Staphylococci can occur very early in colonization even without apparent barrier breech.

Our screening for inflammatory signals in response to Staphylococcal colonization revealed an increased expression of IL-1Β, IL-17A and IL-12p70 for MRSA. The finding that IL-1β is a dominant skin response to particularly MRSA led us to the intricate localization of LCs throughout the skin epidermis, showing cell projections reaching all the way to the skin surface. Out data suggests a limited ability of these cells to phagocytose Staphylococci nor did we note any intracellular bacteria in LCs *in situ.* Internalization of microbes into LCs has long been suggested as a mechanism for antigen presentation to T cells and has been shown for other microbes, but may not be as essential for Staphylococci(18,53,54). We did however show a clear IL-1β response of these cells to a short bacterial exposure with a strong preference to MRSA. This data fits well with the innovative work from the van Sorge lab which demonstrated how the langerin receptor of human LCs binds the wall teichoic acid of MRSA but not *S. epidermidis*(13). This group also showed a clear difference in the interaction of MRSA with human langerin as opposed to the murine receptor(13). Our work further strengthens these findings to human LCs *in situ* with an IL-1β dominant response to human tissue colonization by MRSA. While others have suggested that a majority of IL-1β is secreted by infiltrating neutrophils(10), this cannot be the case in our *ex vivo* model, as no infiltrating immune cells are present. Our work does suggest however that IL-1β may already be a dominant inflammatory signal prior to the arrival of inflammatory cells. Another source of IL-1β in skin is potentially keratinocytes. Similar to our experiments with LC’s, others have shown IL-1β expression from keratinocytes following infection with *S. aureus* but not *S. epidermidis*(55). However, in that work only *S. aureus* was found to move intracellular, not *S. epidermidis* as we found(55).

Taken together, our study highlights the role of the stratum corneum in skin colonization by Staphylococci. Once this tight barrier is breached, tissue resident immune cells are important in the early identification and response to different species, differentiating commensals from pathogens. Further evaluation of the critical role of epidermal LCs in the context of staphylococcal human skin colonization may help reveal whether this is a key pathway to generating skin immunological tolerance to commensal vs potentially pathogenic bacteria.

## Material and Methods

### Ethics statement

Healthy human skin was obtained as surgical excess from elective plastic surgery. Written informed consent was obtained and all procedures were performed according to Swedish National guidelines and approved by the Regionala etikprövningsnämd I Stockholm (ethical approval no: 2015/432-31 and 2023-00567-02).

### Cell lines

The human epidermal keratinocyte cell line (HEKa) (Gibco, Thermo Fischer Scientific Cat# C0055C) was cultured in EpiLife medium (Gibco, Thermo Fischer Scientific Cat# MEPI500CA) supplemented with human keratinocyte growth supplement (Gibco, Thermo Fischer Scientific Cat# S0015) at 37°C with 5% CO2. The human dermal fibroblasts cell line (HDFn) (Gibco, Thermo Fischer Scientific Cat# C0045C) was cultured in Dulbecco’s Modified Eagle Medium/Nutrient Mixture F-12 (DMEM/F-12) (Gibco, Thermo Fischer Scientific Cat# 31330038) supplemented with L-glutamine (Thermo Fischer Scientific Cat# A2916801), 15 mM HEPES and 5% charcoal-stripped fetal bovine serum (FBS) (Gibco, Thermo Fischer Scientific Cat# 12676029) at 37°C with 5% CO2. The MUTZ-3 cell line (DSMZ, no ACC 295) was cultured in Minimum Essential Medium Alpha (MEMα) (Gibco, Thermo Fischer Scientific Cat# 22571020) supplemented with 20% charcoal-stripped FBS, 10% medium conditioned by the 5637 cell line (ATCC no. HTB-9), 1x GlutaMAX supplement (Gibco, Thermo Fischer Scientific Cat# A1286001) and 1x antibiotic-antimycotic (Gibco, Thermo Fischer Scientific Cat# 15240096) at 37°C with 5% CO2. All cells were handled and maintained according to the supplier recommendations and following standard cell culture procedures.

### Bacteria

The bacterial strains used in this study are methicillin resistant *S. aureus* SF8300 USA300-0114(19), further called *S. aureus* USA300 or MRSA, methicillin resistant *S. aureus* USA300-0114 pCM29 (Cam (R), Psar-A sGFP)(56), further called *S. aureus* USA300 GFP, and *S. epidermidis* 1457(20). For all experiments, bacteria were grown in Tryptic Soy Broth (TSB) (Sigma) at 37°C with 180 rpm shaking or on Tryptic Soy Agar (TSA) plates (Sigma) at 37°C overnight. *S. aureus* USA300 GFP was grown in presence of 10µg/ml chloramphenicol. For infections or inoculations, the bacteria were picked from a colony and grown overnight. On the day of the experiment, a fresh culture was re-inoculated and grown to a density of OD600 = 0.3. Bacteria were washed twice in 1xPBS before use. Alternatively, frozen stock bacterial suspensions of ≈1×10^9^ CFU/ml in PBS were used.

### Human skin tissue

Healthy human skin was obtained as surgical excess from elective plastic surgery. Written informed consent was obtained from all participants, and all procedures were approved by the Regionala etikprövningsnämd I Stockholm (approval no: 2015/432-31 and 2023-00567-02). Donors underwent standard surgical preparation which included topical decontamination of the skin with Chlorhexidine. Skin was stored at 4°C until use and cleaned again with 70% ethanol before biopsy preparation.

### *Ex vivo* human skin culture and bacterial colonization

The explant human skin model was prepared and cultured as previously described by Schulz et al. 2019. Briefly, 10 mm full thickness skin biopsies were prepared with a biopsy punch (Agnthos) and placed into a 12 well transwell insert (Sarstedt), supported with CO2 independent media (Gibco, Thermo Fischer Scientific Cat# 18045088) supplemented with 10% FBS and 1% GlutaMax^TM^ (Thermo Fisher Scientific). To seal the edges of the explants, hydrogel rings were prepared from a hydrogel band aid using 11- and 7-mm punches and attached to the top of the explants. Overnight the prepared biopsies stabilized at 37°C in the incubator before topical inoculation with 2 μl of bacteria adjusted to 6.5×10^8^ CFU/ml in 1xPBS (1.3×10^6^ CFU/biopsy). Biopsies treated with 1xPBS were used as negative control. For the *ex vivo* colonization experiment with disrupted stratum corneum, the epidermis of biopsies was carefully scratched with a needle before colonization with bacteria. The applied liquid was allowed to completely evaporate before the explants were incubated at 37°C for 48 h. Culture medium was changed every 24 h. After 48 h the biopsies were homogenized in 500 μl PBS supplemented with 1x HALT protease inhibitor (ThermoScientific, Thermo Fischer Scientific Cat# 87785) using FastPrep lysing matrix A tubes at 6 m/s for 3 x 20 s in a Fast-Prep-24 homogenizer (MP Biomedicals). Serial dilutions of the homogenates were plated on TSA and incubated at 37°C to determine viable CFUs. The remaining skin homogenates were centrifuged at 16000 x g and collected supernatants were stored at -20°C for further analysis. Some biopsies were fixed overnight with 4% paraformaldehyde at 4°C, washed with PBS and used for staining and imaging purposes.

### Dermal/Epidermal sheet separation

To separate the epidermal sheet from the dermis of *ex vivo* human skin biopsies, the samples were incubated in dispase II (Gibco, Thermo Fischer Scientific Cat# 17105041) diluted to 1.8U/ml in CO2 independent medium supplemented with 10% FBS at 4°C overnight. Using sterile tweezers, the epidermis was carefully peeled off the dermis the following day and used for microscopic analysis as described below.

### HEKa and HDFn bacterial infection

Cells were grown to 80% confluency, enzymatically harvested, and seeded in a 48-well plate (Corning) at a density of 2×10^5^ cells/ml for HEKa and 1.6×10^5^ cells/ml for HDFn in 250μl appropriate cell culture media. The following day, the attached cells were infected with the different strains at a multiplicity of infection (MOI) of 1:100. Uninfected cells were used as negative control. After 3 h incubation at 37°C and 5% CO2, the extracellular bacteria were removed in all samples by adding 250μl appropriate cell culture medium supplemented with 100 μg/ml gentamicin and 5 μg/ml lysostaphin (Sigma-Aldrich Cat# L7386) (gentamicin-lysostaphin protection assay) for 30 min at 37°C and 5% CO2. The 3 h post infection (p.i.) samples were washed with 1xPBS, lysed with 0.1% Triton X-100 and serial dilutions were plated on TSA to determine intracellular colony forming units (CFUs). 21 h after removing extracellular bacteria, the supernatants from the remaining samples were collected, centrifuged at 9400 x g for 15 min and stored at -20°C for further analysis.

### MUTZ-3 bacterial infection

To differentiate MUTZ-3 cells into a Langerhans cell phenotype (MUTZ-LC) 2×10^5^ cells/ml MUTZ-3 were exposed to 100ng/ml GM-CSF (PeproTech Cat# 300-03), 10ng/ml TGFβ (PeproTech Cat# 100-21) and 2.5ng/ml TNFα (PeproTech Cat# 300-01A) for 9-10 days in a 12-well plate (1 ml/well). The media was change on day 4 and 8 by and antibiotic-antimycotic was removed on day 8. For the infection assay, MUTZ-LCs were seeded at a density of 2×10^5^ cells/ml in Iscove’s Modified Dulbecco’s Medium (IMDM) (Gibco, Thermo Fischer Scientific Cat# 12440053) supplemented with 10% charcoal-stripped FBS in a 24-well plate (500 µl/well) and infected with *S. aureus* USA300 or *S. epidermidis* 1457 at different MOIs. As for HEKa and HDFn, cell culture supernatants were collected at 3 h p.i. and the infection was stopped using 100 μg/ml gentamicin and 5 μg/ml lysostaphin for 30 min. 24 h p.i. the supernatants were collected again. All supernatants were stored at -20°C for cytokine detection. In addition, intracellular CFUs were determined in cell lysates at 3 h p.i. by plating serial dilutions on TSA.

### Primary corneocyte sampling and bacterial inoculation

After thorough cleaning human skin tissue with 70% ethanol, primary corneocytes were collected using D100 D-squame sampling discs (Clinical & Derm, Thermo Fischer Scientific Cat# NC9714282). Circles with Ø10 mm were punched out and placed into a 48-well plate (Corning). The attached corneocytes were infected with 250μl bacterial suspension at 2×10^7^ CFU/ml in CO2 independent media supplemented with 10% FBS and 1% GlutaMax™ (Thermo Fisher Scientific). After 3h incubation at 37°C and 5% CO2, the supernatant was collected, and extracellular bacteria were removed by adding 250μl medium supplemented with 100 μg/ml gentamicin and at least 5 μg/ml lysostaphin (gentamicin-lysostaphin protection assay). After 30 min incubation, cells were washed with 1xPBS and intracellular CFUs were determined by lysing with 0.3% Triton x-100 for 10 min and plating serial dilutions on TSA. The collected supernatants were centrifuged at 9400 x g for 15 min and stored at -20°C for further analysis. For microscopy, some infected corneocytes were fixed in 4% PFA for 5 min at RT before staining.

### Corneocyte staining for autofluorescence

To visualize the freshly insolated and infected corneocytes (described above) we used 0.4% trypan blue solution to visualize the red autofluorescence of corneocytes following a protocol by Mills et al 2022(57).

### Cytokine detection

Interleukin-8 (IL-8) concentrations were determined with the IL-8/CXCL8 DuoSet ELISA kit (R&D Systems, Cat# DY208), CXCL12 with the CXCL12/SDF-1 DuoSet ELISA kit (R&D Systems, Cat# DY350) and IL-1β concentrations were determined with the ELISA MAX™ Deluxe Set Human IL-1β (BioLegend, Cat# 437004) in 96-well half area Costar assay plates (Corning) according to manufacturer’s protocol with halved volumes. For IL-8 detection, cell culture supernatants from HEKa and HDFn were used at a 10-fold dilution, human skin homogenate supernatants at a 20-fold dilution, and corneocyte supernatants undiluted. For IL-1β detection, cell culture supernatants were used without dilution. Corrected optical density after colorimetric reaction at OD = 450 nm (corrected with OD = 570 nm) was measured with plate reader Tecan Infinite® M1000 pro and analyzed with Excel version 16.79.1 and GraphPad Prism version 10. The panel of cytokines presented in Fig 5 was measured using the LEGENDplex™ Human Inflammation Panel 1 (13-plex) (BioLegend, Cat# 740809) kit, detecting human IL-1β, IFN-α2, IFN-γ, TNF-α, MCP-1, IL-6, CXCL8 (IL-8), IL-10, IL-12p70, IL-17A, IL-18, IL-23 and IL-33. The assay was performed following manufacturer instructions.

### FACS Analysis of MUTZ-LC

Following differentiation of MUTZ-3 into MUTZ-LCs, the differentiation was validated by the measurement of Langerhans cell specific cell surface markers including langerin, CD1a and HLA-DR. MUTZ-LCs were fixed in 4% PFA for 10 min at RT and washed times in 1xPBS followed by blocking with cold 1xPBs supplemented with 10% FBS. Cells were stained with the primary antibodies anti-hu langerin/CD207 clone 929F3.1 (Novus Biologicals, Cat# DDX0362P-100), anti-hu CD1a clone SK9 (BioLegend, Cat# 344902) or anti-hu HLA-DR clone LN3 (BioLegend, Cat# 327002) and the secondary antibodies Alexa Fluor™ 488 anti-rat IgG (Invitrogen, Thermo Fischer Scientific Cat# A-11006), Alexa Fluor™ 555 anti-mouse IgG (Invitrogen, Thermo Fischer Scientific Cat# A-31570) or Alexa Fluor™ 594 anti-mouse IgG (Invitrogen, Thermo Fischer Scientific Cat# A32744). Cell counts were acquired using a Sony ID7000 spectral cell analyzer, data analyzed using FlowJo 10.9.0 and GraphPad Prism 10.

### Microscopy

Fixed (4% paraformaldehyde) human skin samples were embedded in OCT matrix (Tissutek, Saktura) and 5 or 10 µm sections were sliced with an CryoStar™ NX70 cryostat (Thermo Scientific) and collected on SuperFrost® Plus microscopy slides (Thermo Scientific). For H&E staining, the sections were stained in hematoxylin and erythrosin B according to the manufacturer’s protocol and imaged using the Zeiss Axioplan. For immunohistochemistry, the sections were blocked with 0.1% Triton X-100 and 10% FBS in 1xPBS overnight at 4°C and stained with the following reagents and antibodies: NucBlue™ Fixed Cell ReadyProbes™ Reagent (DAPI) (Invitrogen), Alexa Fluor™ 647 Phalloidin 1:500 (Invitrogen, Thermo Fischer Scientific Cat# A22287), anti-cytokeratin-10 1:400 (Invitrogen, clone DE-K10, Thermo Fischer Scientific Cat# MA5-13705), Alexa Fluor™ 555 anti-mouse IgG 1:1000 (Invitrogen). For imaging of the cell monolayers (Fig 2), cells were handled as described above for infections but were seeded in a 8-well slide chamber (ibidi 190711/6) on a permanox plastic Nunc™ Microscope Slide (Thermo Scientific, Cat# 160005). The cells were fixed with 4% PFA for 5 min at room temperature. Blocking and staining was performed as described above. After fixing the epidermal sheets for 2 h in 4% PFA at 4°C, 0.5% Triton X-100 in TBS was used to permeabilize 45 min at 4°C. The sheets were blocked with 0.1% Triton X-100 and 10% FBS in 1xPBS for 2 h at 37°C and stained with the following reagents and antibodies: NucBlue™ Fixed Cell ReadyProbes™ Reagent (DAPI) (Invitrogen, Thermo Fischer Scientific Cat# R37606), Alexa Fluor™ 647 Phalloidin 1:500 (Invitrogen), anti-CD207/langerin 1:400 (novus Biologicals, 929F3.01), Alexa Fluor™ 488 anti-rat IgG 1:500 (Invitrogen). The prepared microscopy samples were mounted in Dako fluorescence medium (Agilent Technologies) and imaged with a Zeiss LSM980-Airy2 (Fig 1) or a Leica Stellaris 5 X (Fig 2,4) Confocal Laser Scanning Microscope (CLSM). Fixed MUTZ-LCs were stained for nuclei, f-actin and langerin using the same antibodies as described above mounted and imaged. Image analysis was performed with Zeiss Zen lite (Version: 3.99.02000), LAS X Office (Version: 1.4.5 27713) or ImarisViewer (Version 10.1.0).

### Statistical Analysis

Statistical tests used were unpaired t test with Welch’s correction to compare two groups, and one-way or two-way ANOVA with Tukey’s or Šídák’s correction for multiple comparisons. The analysis was performed with GraphPad Prism 10 (GraphPad Software, Inc. La Jolla, CA). Statistical significance was indicated with * (p ≤ 0.05), ** (p ≤ 0.01), *** (p ≤ 0.001) or **** (p ≤ 0.0001) or ns (not significant). Details of statistical tests used and the n numbers for each experiment can be found in the figure legends.

## Acknowledgements

The authors thank Mariam Shahata and Tianqi Zhang for technical assistance and support, Professor Liv Eidsmo for valuable discussions and staff at Nordiska Kliniken for facilitating sample collection. Authors acknowledge support from AIMES – Center for the Advancement of Integrated Medical and Engineering Sciences (www.aimes.se) at Karolinska Institutet (1–249/2019) and KTH Royal Institute of Technology (VF-2019–0110), Getinge AB (4–1599/2018). Additional funding was received from Swedish Research Council (2020–01790), Novo Nordisk Funden (NNF22OC0077593) and the German Academic Exchange Service (DAAD). The funders had no role in study design, data collection and analysis, decision to publish, or preparation of the manuscript. Imaging data was collected at the BIC facility, Karolinska Institutet.

## Author contribution

Conceptualization, J.L. and K.M.; Methodology, J.L. and A.B.; Formal Analysis, J.L., A.B. and K.M.; Investigation, J.L. and A.B.; Resources, K.M, M.E.; Writing – Original Draft, J.L. and K.M., Writing – Review & Editing – all authors; Visualization – J.L.; Supervision, J.L. and K.M.; Project Administration – J.L. and K.M.; Funding Acquisition, K.M.

## Lead contact and Materials Availability

Further information and requests for resources and reagents should be directed to and will be fulfilled by the lead author, Dr. Keira Melican. This study did not generate new unique reagents.

## Supporting Information

**Supplementary Fig S1. Tissue integrity of *ex vivo* model.** H&E staining of human skin cross sections 0 h and 72 h after *ex vivo* culture Scale bar = 50µm.

**Supplementary Fig S2. Disruption of stratum corneum**. Representative microscopy image of disrupted stratum corneum by scratching the surface of human skin biopsies. Immunofluorescent staining of keratinocytes in stratum spinosum and granulosum expressing cytokeratin-10 (orange), f-actin (white), nuclei (blue). Scale bar = 50µm.

**Supplementary Fig S3. Confirmation of MUTZ_LC differentiation**. Histograms of stained (red) MUTZ-LCs expressing langerin (CD207), CD1a and HLA-DR vs unstained MUTZ-LCs (grey) analyzed by FACS.

## References

1. Byrd AL, Belkaid Y, Segre JA. The human skin microbiome. Nat Rev Microbiol. 2018 Mar;16(3):143–55.

2. Cogen AL, Nizet V, Gallo RL. Skin microbiota: a source of disease or defence? Br J Dermatol. 2008 Mar 1;158(3):442–55.

3. Otto M. Staphylococcus colonization of the skin and antimicrobial peptides. Expert Rev Dermatol. 2010 Apr 1;5(2):183–95.

4. O’Gara JP. Into the storm: Chasing the opportunistic pathogen Staphylococcus aureus from skin colonisation to life-threatening infections. Environ Microbiol. 2017;19(10):3823–33.

5. Moran GJ, Krishnadasan A, Gorwitz RJ, Fosheim GE, McDougal LK, Carey RB, et al. Methicillin-resistant S. aureus infections among patients in the emergency department. N Engl J Med. 2006 Aug 17;355(7):666–74.

6. Murray CJL, Ikuta KS, Sharara F, Swetschinski L, Aguilar GR, Gray A, et al. Global burden of bacterial antimicrobial resistance in 2019: a systematic analysis. The Lancet. 2022 Feb 12;399(10325):629–55.

7. WHO publishes list of bacteria for which new antibiotics are urgently needed [Internet]. [cited 2024 Jan 8]. Available from: https://www.who.int/news/item/27-02-2017-who-publishes-list-of-bacteria-for-which-new-antibiotics-are-urgently-needed

8. Cho JS, Pietras EM, Garcia NC, Ramos RI, Farzam DM, Monroe HR, et al. IL-17 is essential for host defense against cutaneous Staphylococcus aureus infection in mice. J Clin Invest. 2010 May;120(5):1762–73.

9. Kim HK, Missiakas D, Schneewind O. Mouse models for infectious diseases caused by Staphylococcus aureus. J Immunol Methods. 2014 Aug 1;410:88–99.

10. Miller LS, Pietras EM, Uricchio LH, Hirano K, Rao S, Lin H, et al. Inflammasome-mediated production of IL-1beta is required for neutrophil recruitment against Staphylococcus aureus in vivo. J Immunol Baltim Md 1950. 2007 Nov 15;179(10):6933–42.

11. Holtfreter S, Radcliff FJ, Grumann D, Read H, Johnson S, Monecke S, et al. Characterization of a Mouse-Adapted Staphylococcus aureus Strain. PLOS ONE. 2013 Sep 2;8(9):e71142.

12. Pasparakis M, Haase I, Nestle FO. Mechanisms regulating skin immunity and inflammation. Nat Rev Immunol. 2014 May;14(5):289–301.

13. van Dalen R, De La Cruz Diaz JS, Rumpret M, Fuchsberger FF, van Teijlingen NH, Hanske J, et al. Langerhans Cells Sense Staphylococcus aureus Wall Teichoic Acid through Langerin To Induce Inflammatory Responses. mBio. 2019 May 14;10(3):10.1128/mbio.00330-19.

14. Spaan AN, Surewaard BGJ, Nijland R, van Strijp JAG. Neutrophils Versus Staphylococcus aureus: A Biological Tug of War. Annu Rev Microbiol. 2013;67(1):629–50.

15. Proctor RA. Challenges for a Universal Staphylococcus aureus Vaccine. Clin Infect Dis. 2012 Apr 15;54(8):1179–86.

16. Parker D. Humanized Mouse Models of Staphylococcus aureus Infection. Front Immunol. 2017 May 4;8:512.

17. Schulz A, Jiang L, de Vor L, Ehrström M, Wermeling F, Eidsmo L, et al. Neutrophil Recruitment to Noninvasive MRSA at the Stratum Corneum of Human Skin Mediates Transient Colonization. Cell Rep. 2019 Oct 29;29(5):1074–1081.e5.

18. Pilkington SM, Ogden S, Eaton LH, Dearman RJ, Kimber I, Griffiths CEM. Lower levels of interleukin-1β gene expression are associated with impaired Langerhans’ cell migration in aged human skin. Immunology. 2018;153(1):60–70.

19. Diep BA, Gill SR, Chang RF, Phan TH, Chen JH, Davidson MG, et al. Complete genome sequence of USA300, an epidemic clone of community-acquired meticillin-resistant Staphylococcus aureus. The Lancet. 2006 Mar 4;367(9512):731–9.

20. Mack D, Siemssen N, Laufs R. Parallel induction by glucose of adherence and a polysaccharide antigen specific for plastic-adherent Staphylococcus epidermidis: evidence for functional relation to intercellular adhesion. Infect Immun. 1992 May 1;60(5):2048–57.

21. Grice EA, Kong HH, Conlan S, Deming CB, Davis J, Young AC, et al. Topographical and Temporal Diversity of the Human Skin Microbiome. Science. 2009 May 29;324(5931):1190–2.

22. Nakamura Y, Oscherwitz J, Cease KB, Chan SM, Muñoz-Planillo R, Hasegawa M, et al. Staphylococcus δ-toxin induces allergic skin disease by activating mast cells. Nature. 2013 Nov;503(7476):397–401.

23. Wanke I, Skabytska Y, Kraft B, Peschel A, Biedermann T, Schittek B. Staphylococcus aureus skin colonization is promoted by barrier disruption and leads to local inflammation. Exp Dermatol. 2013;22(2):153–5.

24. Bitschar K, Staudenmaier L, Klink L, Focken J, Sauer B, Fehrenbacher B, et al. Staphylococcus aureus Skin Colonization Is Enhanced by the Interaction of Neutrophil Extracellular Traps with Keratinocytes. J Invest Dermatol. 2020 May 1;140(5):1054–1065.e4.

25. Lang JC, Seiß EA, Moldovan A, Müsken M, Sauerwein T, Fraunholz M, et al. A Photoconvertible Reporter System for Bacterial Metabolic Activity Reveals That Staphylococcus aureus Enters a Dormant-Like State to Persist within Macrophages. mBio. 2022 Oct 26;13(5):e0231622.

26. Tomic-Canic M, Burgess JL, O’Neill KE, Strbo N, Pastar I. Skin Microbiota and its Interplay with Wound Healing. Am J Clin Dermatol. 2020 Sep;21(Suppl 1):36–43.

27. Pastar I, Sawaya AP, Marjanovic J, Burgess JL, Strbo N, Rivas KE, et al. Intracellular Staphylococcus aureus triggers pyroptosis and contributes to inhibition of healing due to perforin-2 suppression. J Clin Invest. 2021 Dec 15;131(24):e133727.

28. Deng L, Costa F, Blake KJ, Choi S, Chandrabalan A, Yousuf MS, et al. S. aureus drives itch and scratch-induced skin damage through a V8 protease-PAR1 axis. Cell. 2023 Nov 22;186(24):5375–5393.e25.

29. Miller LS, Cho JS. Immunity against Staphylococcus aureus cutaneous infections. Nat Rev Immunol. 2011 Aug;11(8):505–18.

30. Valeri M, Raffatellu M. Cytokines IL-17 and IL-22 in the host response to infection. Napier B, editor. Pathog Dis. 2016 Dec;74(9):ftw111.

31. Kurschus FC, Moos S. IL-17 for therapy. J Dermatol Sci. 2017 Sep 1;87(3):221–7.

32. Chen K, Kolls JK. Interluekin-17A (IL17A). Gene. 2017 May 30;614:8–14.

33. Heim CE, Vidlak D, Scherr TD, Hartman CW, Garvin KL, Kielian T. IL-12 promotes myeloid-derived suppressor cell recruitment and bacterial persistence during Staphylococcus aureus orthopedic implant infection. J Immunol Baltim Md 1950. 2015 Apr 15;194(8):3861–72.

34. Posseme C, Llibre A, Charbit B, Bondet V, Rouilly V, Saint-André V, et al. Early IFNβ secretion determines variable downstream IL-12p70 responses upon TLR4 activation. Cell Rep. 2022 Jun 28;39(13):110989.

35. Hamza T, Barnett JB, Li B. Interleukin 12 a Key Immunoregulatory Cytokine in Infection Applications. Int J Mol Sci. 2010 Feb 26;11(3):789–806.

36. Verma ND, Hall BM, Plain KM, Robinson CM, Boyd R, Tran GT, et al. Interleukin-12 (IL-12p70) Promotes Induction of Highly Potent Th1-Like CD4(+)CD25(+) T Regulatory Cells That Inhibit Allograft Rejection in Unmodified Recipients. Front Immunol. 2014;5:190.

37. Miller LS, O’Connell RM, Gutierrez MA, Pietras EM, Shahangian A, Gross CE, et al. MyD88 Mediates Neutrophil Recruitment Initiated by IL-1R but Not TLR2 Activation in Immunity against Staphylococcus aureus. Immunity. 2006 Jan 1;24(1):79–91.

38. Abe T, Sugano E, Saigo Y, Tamai M. Interleukin-1β and Barrier Function of Retinal Pigment Epithelial Cells (ARPE-19): Aberrant Expression of Junctional Complex Molecules. Invest Ophthalmol Vis Sci. 2003 Sep 1;44(9):4097–104.

39. Nakamura Y, Kambe N, Saito M, Nishikomori R, Kim YG, Murakami M, et al. Mast cells mediate neutrophil recruitment and vascular leakage through the NLRP3 inflammasome in histamine-independent urticaria. J Exp Med. 2009 Apr 13;206(5):1037–46.

40. Matsushima H, Ogawa Y, Miyazaki T, Tanaka H, Nishibu A, Takashima A. Intravital Imaging of IL-1β Production in Skin. J Invest Dermatol. 2010 Jun 1;130(6):1571–80.

41. Igyártó BZ, Haley K, Ortner D, Bobr A, Gerami-Nejad M, Edelson BT, et al. Skin-Resident Murine Dendritic Cell Subsets Promote Distinct and Opposing Antigen-Specific T Helper Cell Responses. Immunity. 2011 Aug 26;35(2):260–72.

42. McDermott R, Bausinger H, Fricker D, Spehner D, Proamer F, Lipsker D, et al. Reproduction of Langerin/CD207 Traffic and Birbeck Granule Formation in a Human Cell Line Model. J Invest Dermatol. 2004 Jul 1;123(1):72–7.

43. van den Berg LM, Ribeiro CMS, Zijlstra-Willems EM, de Witte L, Fluitsma D, Tigchelaar W, et al. Caveolin-1 mediated uptake via langerin restricts HIV-1 infection in human Langerhans cells. Retrovirology. 2014 Dec 31;11(1):123.

44. Al Kindi A, Alkahtani AM, Nalubega M, El-Chami C, O’Neill C, Arkwright PD, et al. Staphylococcus aureus Internalized by Skin Keratinocytes Evade Antibiotic Killing. Front Microbiol. 2019 Sep 24;10:2242.

45. Ngo QV, Faass L, Sähr A, Hildebrand D, Eigenbrod T, Heeg K, et al. Inflammatory Response Against Staphylococcus aureus via Intracellular Sensing of Nucleic Acids in Keratinocytes. Front Immunol. 2022 Feb 24;13:828626.

46. McCormack RM, de Armas LR, Shiratsuchi M, Fiorentino DG, Olsson ML, Lichtenheld MG, et al. Perforin-2 is essential for intracellular defense of parenchymal cells and phagocytes against pathogenic bacteria. Cossart P, editor. eLife. 2015 Sep 24;4:e06508.

47. McCormack R, de Armas LR, Shiratsuchi M, Ramos JE, Podack ER. Inhibition of Intracellular Bacterial Replication in Fibroblasts Is Dependent on the Perforin-Like Protein (Perforin-2) Encoded by Macrophage-Expressed Gene 1. J Innate Immun. 2012 Dec 15;5(2):185–94.

48. Barbarot S, Auziere S, Gadkari A, Girolomoni G, Puig L, Simpson EL, et al. Epidemiology of atopic dermatitis in adults: Results from an international survey. Allergy. 2018;73(6):1284–93.

49. Miller LG, Eells SJ, David MZ, Ortiz N, Taylor AR, Kumar N, et al. Staphylococcus aureus Skin Infection Recurrences Among Household Members: An Examination of Host, Behavioral, and Pathogen-Level Predictors. Clin Infect Dis. 2015 Mar 1;60(5):753–63.

50. Kalan LR, Meisel JS, Loesche MA, Horwinski J, Soaita I, Chen X, et al. Strain- and Species-Level Variation in the Microbiome of Diabetic Wounds Is Associated with Clinical Outcomes and Therapeutic Efficacy. Cell Host Microbe. 2019 May;25(5):641–655.e5.

51. Ramirez HA, Pastar I, Jozic I, Stojadinovic O, Stone RC, Ojeh N, et al. Staphylococcus aureus Triggers Induction of miR-15B-5P to Diminish DNA Repair and Deregulate Inflammatory Response in Diabetic Foot Ulcers. J Invest Dermatol. 2018 May 1;138(5):1187–96.

52. Wolcott RD, Hanson JD, Rees EJ, Koenig LD, Phillips CD, Wolcott RA, et al. Analysis of the chronic wound microbiota of 2,963 patients by 16S rDNA pyrosequencing. Wound Repair Regen. 2016;24(1):163–74.

53. Da Silva JP, Da Silva MB, Salgado UI, Diniz JAP, Rozental S, Salgado CG. Phagocytosis of Fonsecaea pedrosoi conidia, but not sclerotic cells caused by Langerhans cells, inhibits CD40 and B7-2 expression. FEMS Immunol Med Microbiol. 2007 Jun 1;50(1):104–11.

54. Vine EE, Rhodes JW, Warner van Dijk FA, Byrne SN, Bertram KM, Cunningham AL, et al. HIV transmitting mononuclear phagocytes; integrating the old and new. Mucosal Immunol. 2022 Apr;15(4):542–50.

55. Vaher H, Kingo K, Kolberg P, Pook M, Raam L, Laanesoo A, et al. Skin Colonization with S. aureus Can Lead to Increased NLRP1 Inflammasome Activation in Patients with Atopic Dermatitis. J Invest Dermatol. 2023 Jul 1;143(7):1268–1278.e8.

56. Pang YY, Schwartz J, Thoendel M, Ackermann LW, Horswill AR, Nauseef WM. agr-Dependent Interactions of Staphylococcus aureus USA300 with Human Polymorphonuclear Neutrophils. J Innate Immun. 2010;2(6):546–59.

57. Mills KB, Roy P, Kwiecinski JM, Fey PD, Horswill AR. Staphylococcal Corneocyte Adhesion: Assay Optimization and Roles of Aap and SasG Adhesins in the Establishment of Healthy Skin Colonization. Microbiol Spectr. 2022 Dec 21;10(6):e0246922.

